# Effects of connectivity hyperalignment (CHA) on estimated brain network properties: from coarse-scale to fine-scale

**DOI:** 10.1101/2024.08.27.609817

**Authors:** Farzad V. Farahani, Mary Beth Nebel, Tor D. Wager, Martin A. Lindquist

## Abstract

Recent gains in functional magnetic resonance imaging (fMRI) studies have been driven by increasingly sophisticated statistical and computational techniques and the ability to capture brain data at finer spatial and temporal resolution. These advances allow researchers to develop population-level models of the functional brain representations underlying behavior, performance, clinical status, and prognosis. However, even following conventional preprocessing pipelines, considerable inter-individual disparities in functional localization persist, posing a hurdle to performing compelling population-level inference. Persistent misalignment in functional topography after registration and spatial normalization will reduce power in developing predictive models and biomarkers, reduce the specificity of estimated brain responses and patterns, and provide misleading results on local neural representations and individual differences. This study aims to determine how connectivity hyperalignment (CHA)—an analytic approach for handling functional misalignment—can change estimated functional brain network topologies at various spatial scales from the coarsest set of parcels down to the vertex-level scale. The findings highlight the role of CHA in improving inter-subject similarities, while retaining individual-specific information and idiosyncrasies at finer spatial granularities. This highlights the potential for fine-grained connectivity analysis using this approach to reveal previously unexplored facets of brain structure and function.

## 1. Introduction

Functional magnetic resonance imaging (fMRI) has provided valuable insights into the neurophysiological underpinnings of human behavior, but the translational potential of fMRI research has yet to be realized. This is partly due to the violation of a key assumption of conventional methods for performing population-level inference in fMRI studies, namely that features (e.g., voxels, vertices, or nodes within a graph representing the brain) are functionally aligned across different individual brains. Large inter-individual differences in functional localization, especially at high spatial resolutions (Haxby et al., 2014), are known to remain after conventional anatomical alignment to a standard stereotaxic space. One area that is particularly vulnerable to this issue is multivariate pattern analysis. Hence, many multivariate models are limited to being subject-specific (Kamitani and Tong, 2005; Norman et al., 2006; Kragel et al., 2018; Al-Wasity et al., 2020). The functional misalignment issue can be partially remedied by spatial smoothing across coarse-grained parcellations (Chen et al., 2017), but at a cost in spatial resolution, effect sizes, and (consequently) power, as high-signal voxels are averaged with low-signal voxels within individuals (a “partial volume” effect). In addition, any remaining misalignment in functional topography will degrade downstream analyses aimed at developing predictive models and biomarkers of behavior/health, and may result in artifactual representations of individual or subgroup differences. This misalignment will affect many measures and outcome metrics, including predictive accuracy, representational similarity, inter-subject correlation (ISC), and graph topological properties, casting doubt on research findings.

Hyperalignment (HA) is a recent analytic approach developed to resolve this issue (Haxby et al., 2011, 2014, 2020). HA is one type in a growing family of “functional alignment” techniques that attempt to register individual brains based on functional properties rather than anatomical locations. Other approaches include the Shared Response Model (Chen et al., 2015) and approaches based on Group Independent Component Analysis (ICA; Du et al., 2020). All these approaches aim to harness variations in activation and/or functional connectivity to create a common functional space. Once (i) a participant’s data are mapped into the functional space based on training data (often, a movie or resting-state scan), and (ii) a predictive model has been developed in the functional space, predictions about the person’s perceptions, feelings, and behavior can be made based on the normative model. Different functional alignment methods make different assumptions about the underlying signal. HA, in particular, assumes that which voxels are active when viewing a particular stimulus are arbitrary and idiosyncratic (i.e., vary arbitrarily across individuals), but that the ‘’representational geometry”—the similarity across multivoxel pattern responses to different stimuli—is preserved across participants. For example, neural patterns when seeing a robin and a parrot may be similar (they are both birds), and both may be dissimilar from those elicited by seeing a monkey. Thus, in HA, brain data from local regions are iteratively mapped into a common high-dimensional space using a Procrustes transformation (Schönemann, 1966), which preserves and aligns participants based on local representational geometry. This procedure increases functional similarities across subjects while preserving idiosyncratic (or subject-specific) information (Haxby et al., 2011; Guntupalli et al., 2016; Feilong et al., 2018; Nastase et al., 2019). The original HA algorithms were “response-based,” aligning the representational geometry of stimulus-evoked responses to watching movies (Guntupalli et al., 2016). More recently, “connectivity-based” approaches (CHA) have been developed that use Procrustes to align the geometry of patterns of correlations with distant brain regions (Guntupalli et al., 2018). Response-based and connectivity-based approaches have also been combined (Busch et al., 2021). Here, we use connectivity-based CHA to functionally align fine-grained cortical connectivity patterns.

Region-of-interest (ROI) CHA (rCHA) and searchlight CHA (sCHA) are two distinct approaches used to perform connectivity-based HA, with the common goal of aligning brain activity patterns across individuals. In rCHA, specific regions of interest are delineated based on anatomical or functional criteria, leveraging prior knowledge of their functional significance or involvement in cognitive processes. Conversely, sCHA operates without predefined ROIs, focusing on fine-grained activity patterns within small, spatially localized regions known as “searchlights,” typically centered around each voxel in the brain. The HA process iterates over all searchlights in the brain, aligning activity patterns within each searchlight across individuals. While rCHA offers computational efficiency and targeted analysis of specific brain regions or networks, sCHA allows for flexible alignment across the entire brain, albeit with increased computational demands. The selection between these methods hinges on the research objectives, available computational resources, and the desired level of spatial specificity for the analysis.

The benefits of functional alignment on functional network measures, including graph theoretic properties and network topology metrics, have yet been explored. That is the focus of the present work. In particular, we seek to explore whether CHA (both rCHA and sCHA techniques) helps improve the performance of graph theoretical analysis of functional brain networks at both coarse and fine spatial scales. A burgeoning literature treats the brain as a multi-scale network in the sense that its connectome—i.e., the map of neural connections—can be modeled over multiple spatial, temporal, and topological scales (Bassett and Siebenhühner, 2013; Betzel and Bassett, 2017). Nearly all studies of “connectomics”, which attempt to map functional networks and their properties, rely on anatomical alignment, assuming that brain voxels or “parcels” (i.e., groups of voxels that form a contiguous and presumably homogenous region, discussed below) located in the same anatomical locations are functionally isomorphic and comparable across individuals. Some procedures have been developed to “individualize” the locations of functional regions based on connectivity, which can partially relax this assumption (Robinson et al., 2014; Glasser et al., 2016; Schaefer et al., 2018). However, the substantial benefits of HA in increasing inter-subject alignment and stimulus classification (Haxby et al., 2011, 2020) suggest that this assumption is typically violated, and different network organization and properties might emerge if HA were used to define a functional reference space before network analysis.

In addition to revealing new principles of network organization, a benefit of HA may be preserving functional information at fine spatial scales. The spatial scale concerns the granularity of brain architecture, ranging from the cellular microscale level of neurons and synapses (Jarrell et al., 2012; Shimono and Beggs, 2015; Schröter et al., 2017; Zheng et al., 2018; Reimann et al., 2019; Cook et al., 2020; Kajiwara et al., 2021) to the macroscale level of regions, pathways, and the whole brain (Bullmore and Bassett, 2011; Craddock et al., 2013). The analysis of human brain networks using fMRI is restricted by the spatial granularity of the individual voxel (or vertex), and cellular level activation is not inherently measurable using fMRI. A typical fMRI voxel is around 3 mm^3^ in size, which can be reduced to 500 μm^3^ or less by increasing the magnetic field (Shmuel et al., 2007) and/or using modern acquisition techniques (e.g., multiband; Feinberg and Setsompop, 2013). The spatial scale captured by high-resolution, single-participant activity patterns contains a great deal of information about stimuli, functional states, and behaviors (Haxby, 2012).

Although voxel-level granularity is considered the lower bound of the MRI spatial scale, it is not commonly used in the analysis of human brain networks due to functional and anatomical variations across individuals, noise effects, signal dropout, and computational cost over large number of voxels (Betzel and Bassett, 2017). To overcome these issues, voxels are typically grouped together into a set of non-overlapping parcels or regions of interest (ROIs), so that brain networks can be constructed based on the averaged characteristics of parcels (de Reus and van den Heuvel, 2013). Atlases and parcellations can be based on resting-state functional connectivity MRI (Power et al., 2011; Thomas Yeo et al., 2011; Gordon et al., 2016; Schaefer et al., 2018), automated anatomical templates (Tzourio-Mazoyer et al., 2002; Destrieux et al., 2010), diffusion MRI (Lefranc et al., 2016), or integrated information from multiple modalities such as myeloarchitecture, resting-state, and task-based fMRI (Glasser et al., 2016; Ji et al., 2019). However, parcellation techniques have major drawbacks related to voxel aggregation and averaging information at these coarse scales. Simply put, any given parcel is likely to average over distinct, fine-grained patterns that relate to distinct functions and networks (Woo et al., 2014). Another issue relates to the reproducibility of examining network topology impacted by the choice of parcellation and resolution (Fornito et al., 2010; Zalesky et al., 2010). HA may ameliorate some of these drawbacks and better align networks with fine-grained pattern information.

The main goal of the paper is to evaluate the effect of CHA on graph analysis using (i) coarse-grained (i.e., brain parcels) and (ii) fine-grained (i.e., individual voxels/vertices) brain data. We used the fMRI data of 200 subjects from the Human Connectome Project (HCP). The HCP is ideal for this analysis due to the relatively large amount of high-quality resting-state fMRI data available for each participant, which is critical to reliably estimating participant-specific functional connectivity patterns, while including enough participants for meaningful characterization at the population level. In the first phase, we used the timeseries of the first run (REST1_LR) to create both ROI-based and searchlight CHA models (a group functional reference space) and tune its parameters. Then, we functionally aligned the timeseries from the other three runs (REST1_RL, REST2_LR, and REST2_RL) to the models. In the second phase, we examined whether CHA led to graph topological changes in the brain organization at coarse-grained and/or fine-grained scales, and if so, whether it helps predict an external variable such as fluid intelligence. Cross-validated prediction (i.e., brain decoding) accuracy is increasingly being used as a benchmark for the strength of brain-stimulus, brain-behavior, and brain-mind outcomes more generally. Here we perform this benchmarking by evaluating whether CHA increases prediction accuracy when predicting a cognitive variable. To this end, we used a series of widely used features in graph theory to predict fluid intelligence. An overview of our analysis pipeline is shown in **Figure 1**.

**Figure 1.**
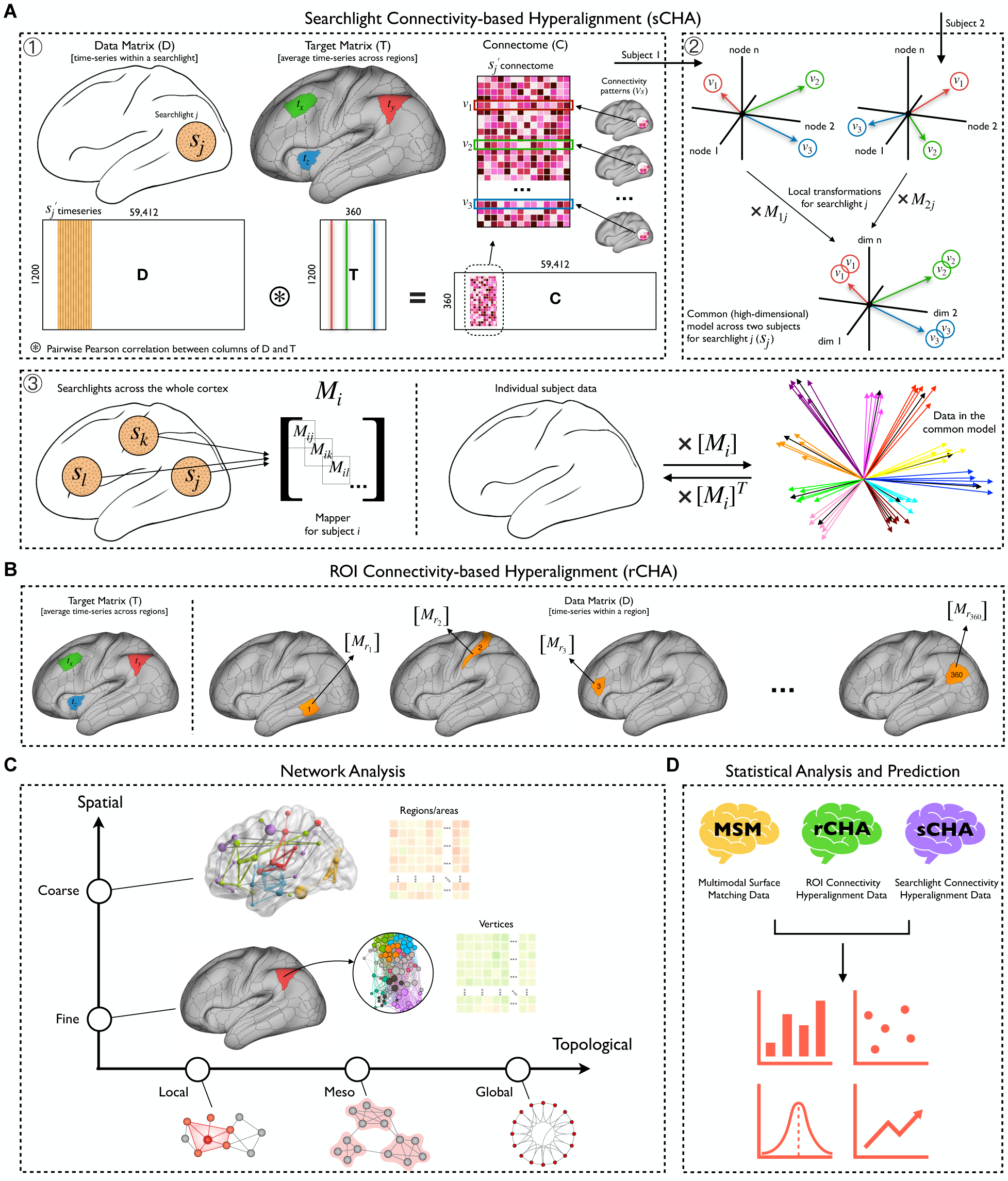
An overview of the analysis pipeline. (A) Searchlight connectivity hyperalignment (sCHA) is performed in three steps: first, a connectome is created for each participant (i.e., a similarity matrix made up of data and targets); second, for a given searchlight across participants, the local transformation per subject and the common model is computed (here, an example for two subjects is shown); third, a mapper (whole-cortex transformation) is obtained for each participant by aggregating their local transformations. (B) ROI connectivity hyperalignment (rCHA) is performed independently for each region. The target matrix is similar to sCHA, but the data matrix is vertex time series within each region. Unlike sCHA, there is no mapper aggregation, making rCHA computationally faster. (C) Network analysis in this study encompasses spatial and topological scales, including coarse and fine spatial scales, as well as local, meso, and global topological properties. Connectomes are constructed at each spatial scale using both original and hyperaligned data (multimodal surface matching (MSM) and connectivity hyperalignment (rCHA and sCHA) data, respectively). These connectomes are then utilized to compute networks and common graph measures across local, meso, and global topological scales. (D) Group-level statistical analysis and predictive modeling were performed on the extracted measures.

## 2. Methods

### 2.1. HCP resting-state fMRI data (rs-fMRI)

As a proof of concept, we selected the first 200 participants from the HCP 1200-subjects release for whom all resting-state runs were available (Van Essen et al., 2013). To enhance the robustness of our analysis, we further refined the sample by excluding twins, including both monozygotic and non-monozygotic individuals. For each subject, four 14.4-minute runs of rs-fMRI (1200 frames) were acquired in two separate sessions/days (REST1 and REST2), with scans in both the left-right (LR) and right-left (RL) phase-encoding directions (Smith et al., 2013). Functional images were collected with an isotropic spatial resolution of 2 mm and a temporal resolution of 0.72 s on a 3T Siemens Skyra scanner using a multi-band sequence. The rs-fMRI data underwent the HCP minimal preprocessing pipeline (Glasser et al., 2013, 2016). Functional images were first spatially realigned, distortion corrected, and normalized to the Montreal Neurological Institute (MNI) template. Then both cortical and subcortical data were processed using the ICA-FIX method (Griffanti et al., 2014; Salimi-Khorshidi et al., 2014) and stored in CIFTI format. This procedure included the regression of 24 motion-related parameters (the six classical motion parameters, their derivatives, and the squares of these 12 parameters). After volumetric preprocessing, rs-fMRI data were projected into the 32k fs_LR surface space by the multimodal surface matching method (MSM-All or MSM; Robinson et al., 2014; Van Essen et al., 2013), which is based on multiple descriptors of brain architecture, function and connectivity. In this study we used cortical data with 32,000 ‘grayordinates’ (vertices) per hemisphere. For model training (i.e., obtaining HA and regression parameters), we used data from the first run (REST1_LR) and tested the models on the remaining three runs (REST1_RL, REST2_LR, and REST2_RL).

### 2.2. Connectivity-based hyperalignment (CHA)

In this study, both surface-based searchlight and ROI-based approaches were utilized to perform CHA. While the searchlight method employs a vertex-centric analysis, the ROI approach focuses on predefined brain regions or networks. For both approaches, we initially masked the whole brain CIFTI files (91,282 grayordinates representing the gray matter as cortical surface vertices as well as subcortical voxels) to include only the cortex, resulting in 59,412 nodes across both hemispheres.

We begin by creating the input to both CHA algorithms (rCHA and sCHA), which is a connectivity matrix—hereafter referred to as a connectome—for each subject (see **Figure 1A**, first box). Each participant’s connectome, **C**, is a type of correlation matrix calculated from a data matrix **D** (i.e., rs-fMRI time-series), and a connectivity target matrix **T**. We defined connectivity targets using the Glasser atlas (Glasser et al., 2016), which partitions the whole brain into 360 regions. We used the average time-series in each region as the target. Assuming **D** to be a 1,200 (number of timepoints in each run) x 59,412 (cortical grayordinates) matrix and **T** as a 1,200 x 360 matrix, **C** is the pairwise Pearson correlation between all features of **D** (vertex time-series) and all features of **T** (target or average parcel time-series), resulting in a 360 x 59,412 connectivity matrix. Thus, each row in **C** represents a *connectivity pattern*—a vector of 59,412 correlation coefficients—across all cortical vertices for a predefined target region, and each column in **C** represents a *connectivity profile*—a vector of 360 correlation coefficients—for each of the cortical grayordinates. Each connectivity profile (i.e., each column of a participant’s connectome) was z-scored to have zero-mean and unit variance, and the normalized connectomes were fed into the searchlight hyperalignment algorithm.

#### Searchlight approach (sCHA)

The searchlight hyperalignment algorithm utilizes Procrustes transformations to create a transformation matrix for each participant that maps their connectome into a high-dimensional information space shared across participants (Guntupalli et al., 2016, 2018). The term “searchlight” indicates that the hyperalignment algorithm centers a searchlight (a spherical region) on each cortical vertex and computes a common information space across participants (**Figure 1A**, second box), resulting in a local transformation matrix per searchlight and subject (for each subject, a total of 59,412 transformations are computed). In this study, we adopted a 10 mm searchlight radius for the hyperalignment process, strategically balancing computational efficiency with neuroanatomical relevance, given the substantial participant cohort and the limited observed gains in ISC associated with using larger radii. The use of searchlights not only makes the Procrustes analysis computationally tractable but also limits the functional alignment to a neuroanatomically meaningful radius (Guntupalli et al., 2016; Feilong et al., 2018). Finally, the local transformation matrices are aggregated to form a whole-cortex transformation matrix for each participant (**Figure 1A**, third box), also called a *mapper* (Haxby et al., 2020).

#### ROI approach (rCHA)

In contrast to the searchlight approach, the ROI approach involves selecting specific regions or networks of interest for analysis. This method begins by identifying relevant brain regions based on prior knowledge or functional parcellation atlases; we used the 360 cortical regions from the Glasser atlas. The connectivity patterns within these regions are then analyzed independently and a mapper obtained for each individual per region.

In the current study and for both approaches, we used the mappers obtained from the first run (i.e., REST1_LR, which served as our model training data) for two principal purposes: (1) to project the connectomes derived from distinct runs (i.e., REST1_RL, REST2_LR, and REST2_RL) into the common space to validate the hyperalignment model; and (2) to project the initial data matrix (i.e., rs-fMRI time-series) into the common space for post hoc analyses (e.g., network analysis and prediction). In the validation phase, the inter-subject correlation (ISC) of the connectivity profiles were calculated for mapped test datasets. The ISC is the pair-wise correlation between the connectivity profiles from a certain vertex averaged across all possible pairs of subjects. Higher values reflect reduced inter-subject error variability.

### 2.3. Network construction

#### Coarse-scale

For all participants in both non-hyperaligned and hyperaligned groups, we parcellated the cortex into 360 ROIs using the Glasser atlas (Glasser et al., 2016). Next, the pairwise functional connectivity between ROIs was computed using Pearson’s correlation, resulting in a 360 × 360 fully weighted matrix for each subject. Finally, a set of global and local graph properties were measured for each participant (Rubinov and Sporns, 2010). See **Figure 2** for mathematical definitions and explanations of the measures used in this study. A permutation test (50,000 permutations) was applied to examine the variations of network properties pre and post CHA (*p*-values corrected using the Benjamini-Hochberg (BH) procedure with q=0.05).

**Figure 2.**
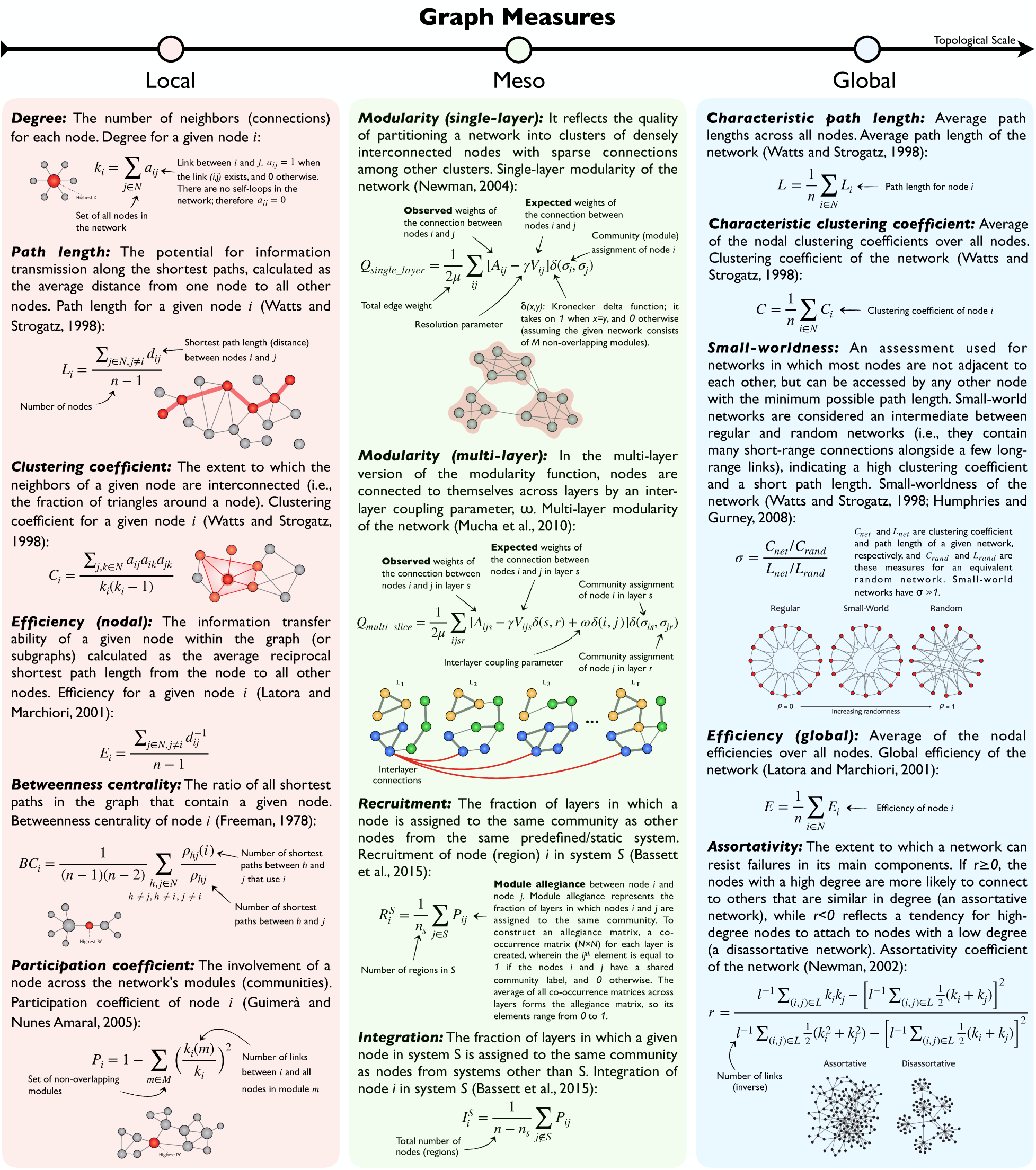
Mathematical definitions, explanations, and visualizations of the various graph measures and metrics used in this study. Based on the topological scale, the measures and metrics are divided into local (pink), meso (green), and global (blue) classes. Here local measures are computed separately for each node, meso measures are computed on systems of nodes, and global measures are computed on the entire graph.

#### Fine-scale

For all participants in both non-hyperaligned and hyperaligned groups, we first projected the initial rs-fMRI time-series in each run to the common space using the created mappers. Then, the new vertex-level time-series were extracted for all 360 Glasser ROIs separately for all subjects. The number of voxels in the ROIs ranges from 31 to 839 (∼165 on average). Then, functional connectivity matrices at fine-scale were constructed separately for each ROI using Pearson correlation. Here fully weighted networks were used with the main diagonal (self-connections) set to 0. Next, global and local graph theory criteria were calculated on the obtained matrices.

To perform meso-scale analysis, we replicated this process using Yeo’s 17 networks atlas (Schaefer et al., 2018), chosen for its consistent vertex distribution across networks (ranging from 1307 to 5980 vertices). This choice facilitated a more balanced and less intricate modular analysis compared to the Glasser atlas with Cole/Anticevic network labels, which exhibited greater variability in network sizes (ranging from 718 to 11537 vertices). Subsequently, we extracted meso-level features (module allegiance, integration, recruitment coefficients) using the generated functional connectivity matrices, and assessed community structures before and after CHA. Further details are outlined below.

### 2.4. Graph metrics calculation

Utilizing the NetworkX library in Python, we investigated the topological properties of functional brain networks (binary undirected matrices) for each participant at global and local levels. A density-based thresholding was used to maintain the strongest links. The mathematical definition of each network statistic is listed in **definitions,** explanations, and visualizations of the various graph measures and metrics used in this study. Based on the topological scale, the measures and metrics are divided into local (pink), meso (green), and global (blue) classes. Here local measures are computed separately for each node, meso measures are computed on systems of nodes, and global measures are computed on the entire graph.

. Global metrics (characteristic path length, mean clustering coefficient, global efficiency, assortativity, single-layer modularity, and small-worldness) principally measure the functional segregation and integration of brain networks. Local metrics (node centrality and hub density) measure the regional properties of networks. Hubs can be classified as provincial or connector, whether they mostly contain local connections within a module or connecting nodes in different modules. Degree, eigenvector centrality, closeness centrality, nodal clustering coefficient, k-core score, and participation coefficient were among the most common local properties calculated in the literature (Rubinov and Sporns, 2010).

In our meso-scale analysis, we applied an iterative multi-layer community detection technique (Mucha et al., 2010) to study the modular organization of the large-scale brain networks across subjects at fine spatial resolution. Each layer represents a subject’s connectivity matrix (or, more precisely, its modularity matrix^1^), which is categorically coupled with other layers/subjects using an interlayer parameter, *ω*, to keep community assignments (labels) consistent across layers. The structural resolution *γ*, which tunes the size of the modules within each layer, is another parameter of the multi-layer modularity function (see **Figure 2** for mathematical definitions). To obtain an appropriate value for *γ* and *ω*, we first created a parametric space (*γ* ∈ [0.5, 1.5] with a step size of 0.1; *ω* ∈ [0, 1] with a step size of 0.1) and chose the combination with the highest *Q* across the majority of networks, yielding values of *γ* = 1.0 and *ω* = 0.1. The modularity index *Q* and community labels *C_ij_* (for node *i* in layer *j*) are the two primary byproducts of the modularity maximization function; the former measures partitioning quality, and the latter is utilized to form the *allegiance matrix* and extract mesoscale (intermediate) properties such as *recruitment* and *integration*. The module allegiance matrix reflects the likelihood of two nodes belonging to the same community (Bassett et al., 2015). To create an allegiance matrix, we built a co-occurrence matrix (number of ROI × number of ROI) for each layer, wherein the (*i, j*)*^th^* element equals 1 if nodes *i* and *j* have a shared community label and 0 otherwise. The average of all co-occurrence matrices across layers (200 layers per run) forms the allegiance matrix; thus, its elements range from 0 to 1. Finally, using the allegiance matrix, we estimated two mesoscale coefficients called *recruitment* and *integration* (Bassett et al., 2015) to compare community structure before and after connectivity hyperalignment (see **Figure 2** for mathematical definitions).

### 2.5. Regression models

We used derived graph theory measures as features to predict fluid intelligence, aiming to compare predictive performance on graph features from hyperaligned and non-hyperaligned data. Our approach involved fitting a linear regression model with a least-squares loss function and l2-norm regularization (aka Ridge Regression). Utilizing rs-fMRI data from MSM, rCHA, and sCHA (representing non-hyperaligned, searchlight, and ROI hyperaligned data, respectively), we derived patterns of graph-based measures from the first dataset (REST1_LR) to serve as training data for the prediction model. Subsequently, the model was tested using patterns of graph measures computed from other runs (REST1_RL, REST2_LR, and REST2_RL). For each participant, the pattern for a certain global measure is a vector of 360 elements calculated one-by-one, corresponding to the fine-grained topological information of that measure in 360 brain regions. The pattern for a certain local measure is calculated for each brain region separately, resulting in a vector of length equal to the number of vertices in that region, which ranges from 31 to 839. Therefore, considering a sample size of 200, an input matrix with dimensions 200 x 360 is obtained for each global measure, and 360 input matrices with dimensions 200 x *n* are obtained for each local measure (where *n*, the number of features, varies between 31 and 839). The target variable is a 1d-array of length 200, consisting of fluid intelligence scores for each individual. The use of one run of resting-state data to train the model and the remaining three to test them is based on previous work in the area (Guntupalli et al., 2018).

We evaluated our models using the mean square error (MSE), a risk metric representing the expected value of the squared (quadratic) error or loss. If ŷ*_i_* is the predicted value of the *i*^th^ sample, and y*_i_* is the corresponding true value, then the mean squared error (MSE) estimated over n*_samples_* is defined as:

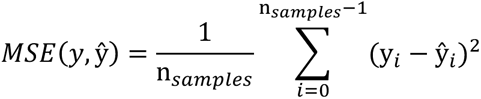

Finally, a paired t-test with Bonferroni-corrected *p*-values was used to compare the MSE values obtained using MSM, rCHA, and sCHA. A chance analysis was also conducted by shuffling the target labels for the training set alongside our regression modeling, evaluating the chance MSE as a baseline to demonstrate that the actual predictive model is capturing meaningful information from the data.

## 3. Results

### 3.1. Connectivity profiles are more similar across participants following searchlight CHA (sCHA)

For each vertex, the inter-subject correlation (ISC) of connectivity profiles was calculated for mapped test sets as the pairwise correlation between profiles averaged across all possible pairs of subjects (**Figure 3A** and **3B**). Clearly, performing sCHA leads to a large increase in ISCs of connectivity profiles in the rs-fMRI data, and the increase is spread throughout the cortex. This increase reflects the ability of the sCHA algorithm to boost the shared signal across individuals and suppress noise. In contrast, performing rCHA does not provide a similar increase in ISC. **Figure 3C** shows the extent to which the mean distribution of ISC values across the cortex changes following hyperalignment. The distribution after sCHA is shifted upward, reflected by the increased mean value of the distribution (0.34 vs. 0.54), while the distribution after rCHA is not significantly shifted (left panel). Finally, a scatter plot (with a linear regression line superimposed) of individual whole cortex mean ISCs of connectivity profiles before and after hyperalignment for both rCHA and sCHA is depicted in **Figure 3D**. Each point represents a subject; the averaged ISC for a vertex is first calculated between a given individual and others, then averaged over the whole cortex.

**Figure 3.**
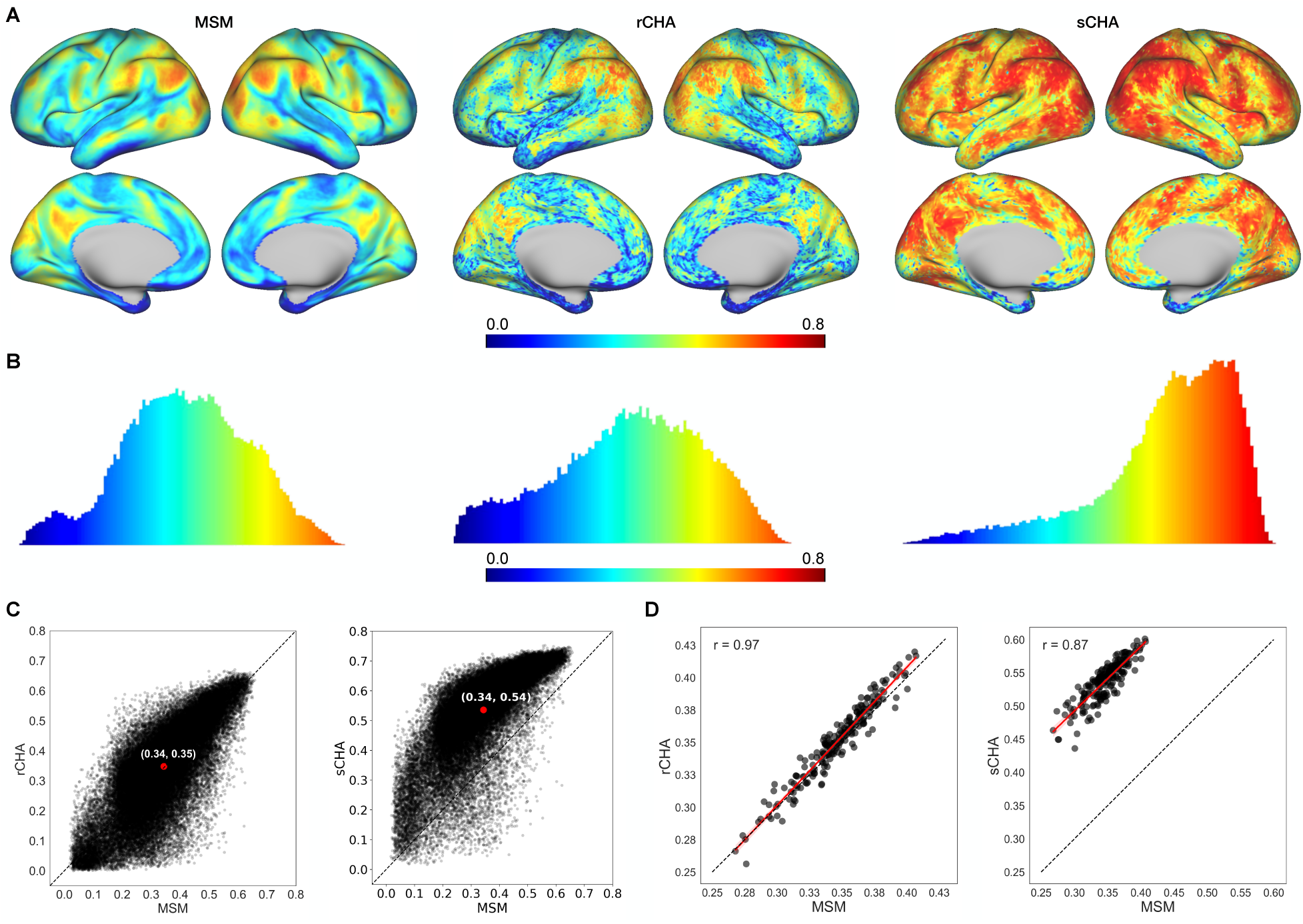
Comparing connectivity profiles pre and post CHA. (A) Mean inter-subject correlations (ISCs) before (MSM) and after hyperalignment (rCHA and sCHA) mapped on the cortical surface, showing increased similarity post hyperalignment, particularly with sCHA. (B) Distribution of ISCs across vertices for MSM, rCHA, and sCHA, with a shift towards higher correlation values post hyperalignment, especially with sCHA. (C) Scatterplots depict ISCs across all vertices before and after CHA. Points above the dashed line (y=x) indicate higher ISCs following hyperalignment. Comparisons between MSM and rCHA (left panel) and MSM and sCHA (right panel) highlight the effectiveness of sCHA in enhancing inter-subject correlation. (D) Scatterplot of individual whole cortex mean ISCs of connectivity profiles pre and post CHA. The left panel compares MSM with rCHA, while the right panel compares MSM with sCHA.

While the ISC results confirm CHA’s capacity (at least sCHA’s capacity) to enhance the shared signal-to-noise ratio across subjects and reduce inter-subject variability in cortical topology, the central query revolves around whether CHA can improve subject-specific outcomes and predictive performance based on topographic idiosyncrasies. To explore this, we extracted a set of graph-based features from both non-hyperaligned and hyperaligned rs-fMRI datasets to evaluate their predictive ability regarding fluid intelligence. This analysis was conducted at two spatial scales—coarse and fine—to assess whether inter-subject variations in cortical architecture become more reliable post-CHA. Investigations were carried out for both rCHA and sCHA conditions for comparative insights.

### 3.2. Parcellation of CHA-aligned data (compared to non-aligned data) reflects the brain’s organization more integrated both locally and globally

Using the estimated mappers, we projected the initial test time-series onto the CHA-derived common space for both techniques (rCHA and sCHA) and performed coarse-scale network analysis before and after CHA, as illustrated in **Figures 4** and **5**. When performing global analysis (**Figure 4**), the findings show a significant decrease in global efficiency following CHA (particularly, sCHA), indicating a shift in the efficiency of information propagation across the network. Concurrently, multiple network attributes notably increased, such as clustering coefficient, assortativity, single-layer modularity, post sCHA (for definitions of the metrics, see **Figure 2**).

**Figure 4.**
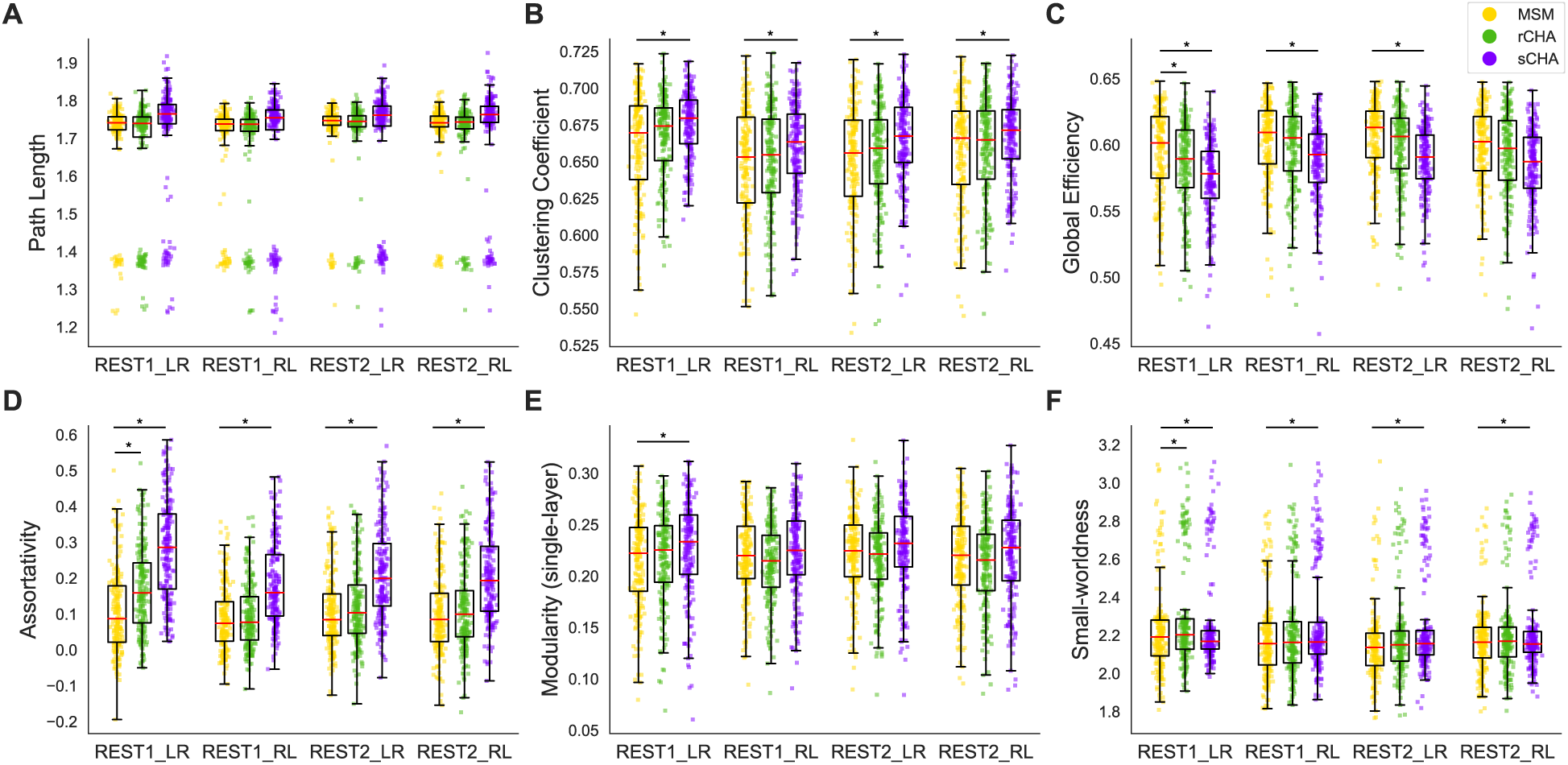
Global properties before/after CHA in coarse-scale structure evaluated in the three test datasets; following hyperalignment, functional brain organization appears more integrated globally. The properties evaluated include (A) path length, (B) clustering coefficient, (C) global efficiency, (D) assortativity, (E) single-layer modularity, and (F) small-worldness. Values for MSM, rCHA, and sCHA are depicted in yellow, green, and purple, respectively. Significant differences in properties before/after CHA are indicated with a * (p-value < 0.05 paired t-test).

**Figure 5.**
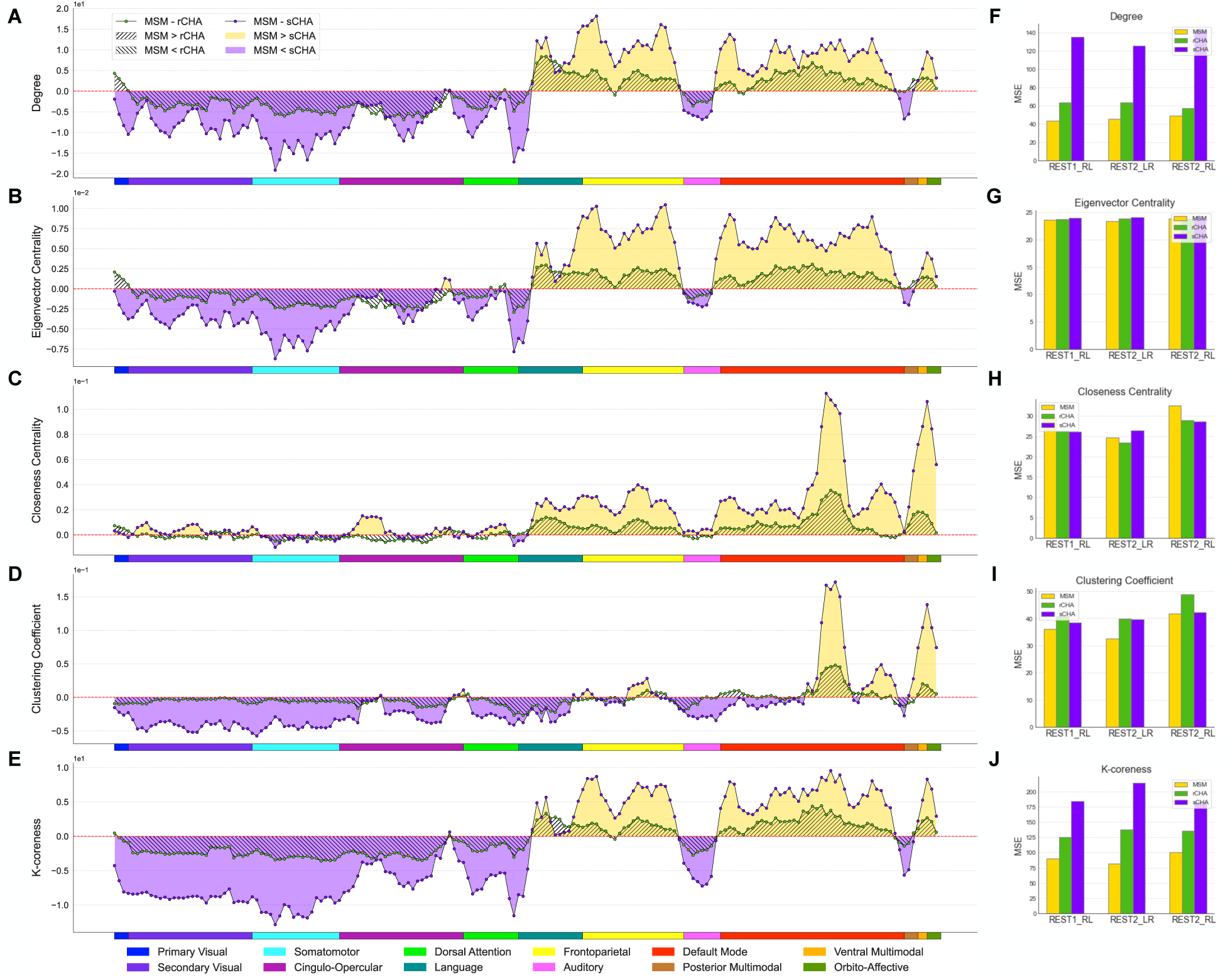
Differences in local properties before/after CHA in coarse-scale structure shown in 360 brain regions grouped into twelve different brain networks. Following CHA (particularly, sCHA), functional brain organization appears more integrated locally but not predictive. The properties evaluated include (A) degree, (B) eigenvector centrality, (C) closeness centrality, (D) clustering coefficient, and (E) k-core score. Values for MSM, rCHA, and sCHA are depicted in yellow, green, and purple, respectively. Panels (F-J) compare MSE before/after CHA in predicting fluid intelligence for each local measure (the feature vector contains 360 elements corresponding to brain regions).

The decreased global efficiency suggests a more localized and interconnected organization in the functionally aligned data. This hints at a different configuration of communication pathways, enhancing information exchange within closely connected brain regions. Conversely, the rise in clustering coefficient, assortativity, modularity, and small-worldness metrics collectively implies increased functional integration among neighboring regions, forming specialized clusters. The seemingly counterintuitive decrease in global efficiency could reflect a trade-off between global information transfer and localized communication hub establishment. This aligns with the brain’s capacity for efficient information processing, using specialized networks for cognitive functions. The concurrent increase in network attributes supports the “small-world” architecture idea, where interconnected modules balance segregation and integration for optimal functionality.

We further investigated local properties (**Figures 5A-E**) and our findings revealed distinct shifts in centrality metrics and nodal clustering coefficients within specific brain networks following the CHA procedure (particularly, sCHA). These changes illuminate a difference in brain hubness and local connectivity patterns. Specifically, we observed enhanced centrality metrics and increased nodal clustering coefficients within networks like the secondary visual, somatomotor, cingulo-opercular, dorsal attention, auditory, and posterior multimodal networks, suggesting increased local integration and information flow efficiency in these networks is observable after hyperalignment. A converse trend emerged within the language, frontoparietal, default, and ventral multimodal networks post-CHA, possibly indicating an alternative allocation of resources or differing interconnectivity dynamics. In each subplot, each point represents the regional value of the corresponding metric across the 360 brain regions; yellow, green, and purple dots correspond to MSM, rCHA, and sCHA, respectively. Twelve different brain networks (taken from the Cole/Anticevic atlas) are also shown in different colors at the bottom of each subplot.

In **Figures 5F-J**, we observed mainly a larger MSE in the prediction of fluid intelligence following coarse-scale analysis. This discrepancy highlights that while CHA enhances measures of network connectivity at coarser levels, it does not yield a corresponding improvement in capturing individual variability, potentially suggesting that the smoothing of hyperaligned fMRI data across coarse-grained parcellations might contribute to the attenuation of unique individual-specific patterns. We also tested the model against chance by shuffling target values in the training set, confirming its superior performance.

### 3.3. CHA improves prediction of fluid intelligence using fine-scale network properties

In this section, we examined the impact of CHA on the prediction of fluid intelligence using fine-scale local graph properties across all three test sets (REST1_RL, REST2_LR, and REST2_RL). As illustrated in **Figure 6A**, a marked reduction in MSE becomes evident in various network connectivity measures following both types of CHA (particularly, rCHA) compared to MSM alignment (verified by paired t-tests with Bonferroni correction). This reduction implies improved predictive capability using CHA-aligned data. The decrease in MSE was observed across all evaluated measures—degree, eigenvalue centrality, closeness centrality, clustering coefficient, and k-coreness—for every test set after CHA. Notably, eigenvalue centrality, closeness centrality, and clustering coefficient emerged as strong predictors of fluid intelligence, outperforming other metrics. We also assessed the model’s performance against chance by randomly shuffling the target values within the training set, revealing that the actual predictive model outperformed chance across all measures. These findings underscore CHA’s role in refining our estimation and modeling of individual brain networks, thereby facilitating a more effective capture of the relationships existing between local graph properties and cognitive traits such as fluid intelligence.

**Figure 6.**
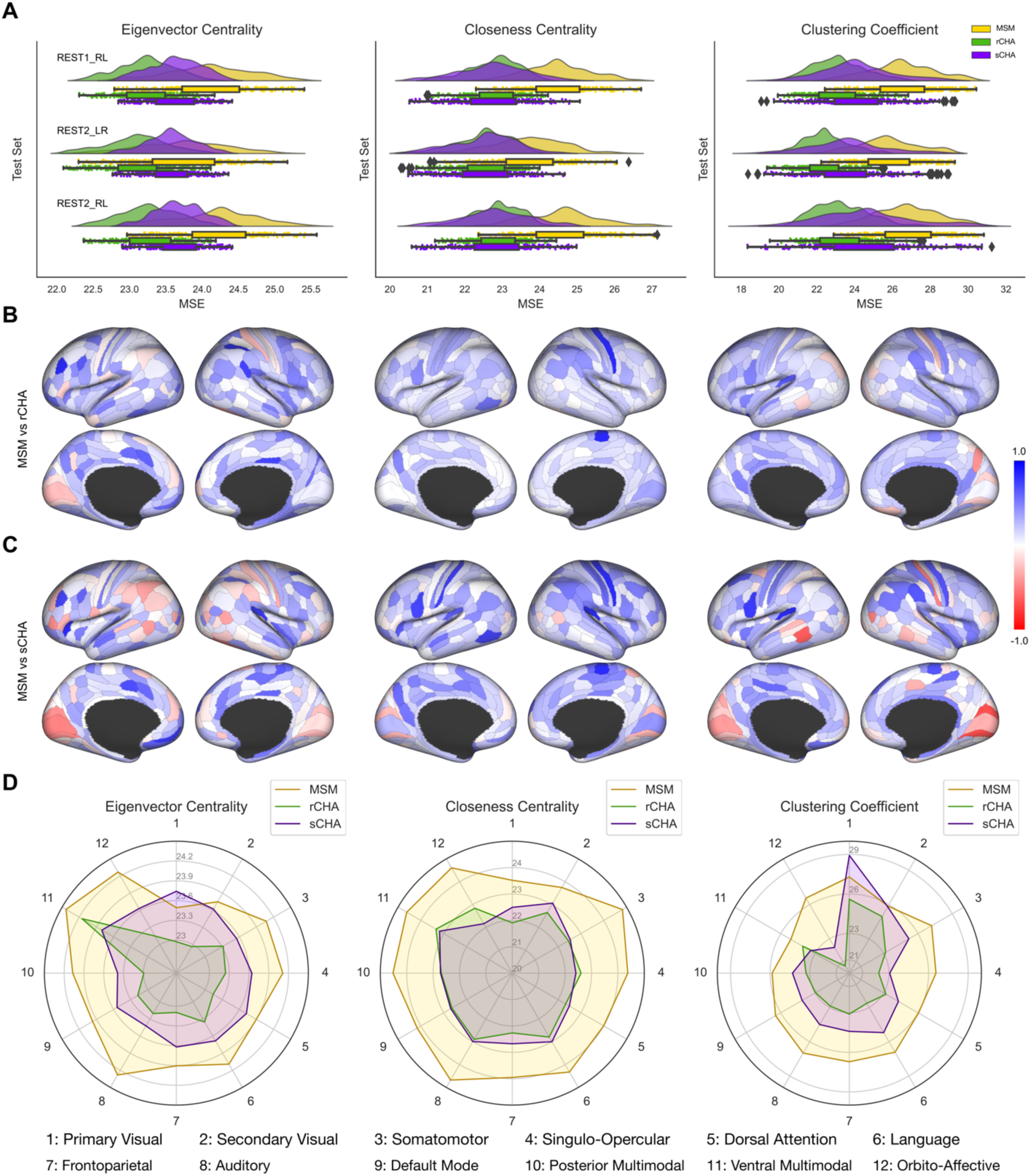
(A) Mean squared error (MSE) of predicting fluid intelligence—calculated separately for each region— based on the pattern of fine-grained local measures derived from MSM, rCHA, and sCHA data series (i.e., rs-fMRI before and after hyperalignment) for the three test datasets. Values for MSM, rCHA, and sCHA are represented in yellow, green, and purple, respectively. (B, C) Normalized mean squared error differences between MSM and rCHA, and MSM and sCHA (i.e., Δ/*max*(*absolute*(Δ)) where Δ= *MSE_MSM_* – *MSE_CHA_*), evaluated for eigenvalue centrality, closeness centrality, and clustering coefficient, encompassing all three test datasets. Positive values indicate improved prediction accuracy after CHA. (D) Average MSE across brain regions, grouped based on the 12 Cole/Anticivic networks shown for eigenvector centrality, closeness centrality, and clustering coefficient.

**Figure 7.**
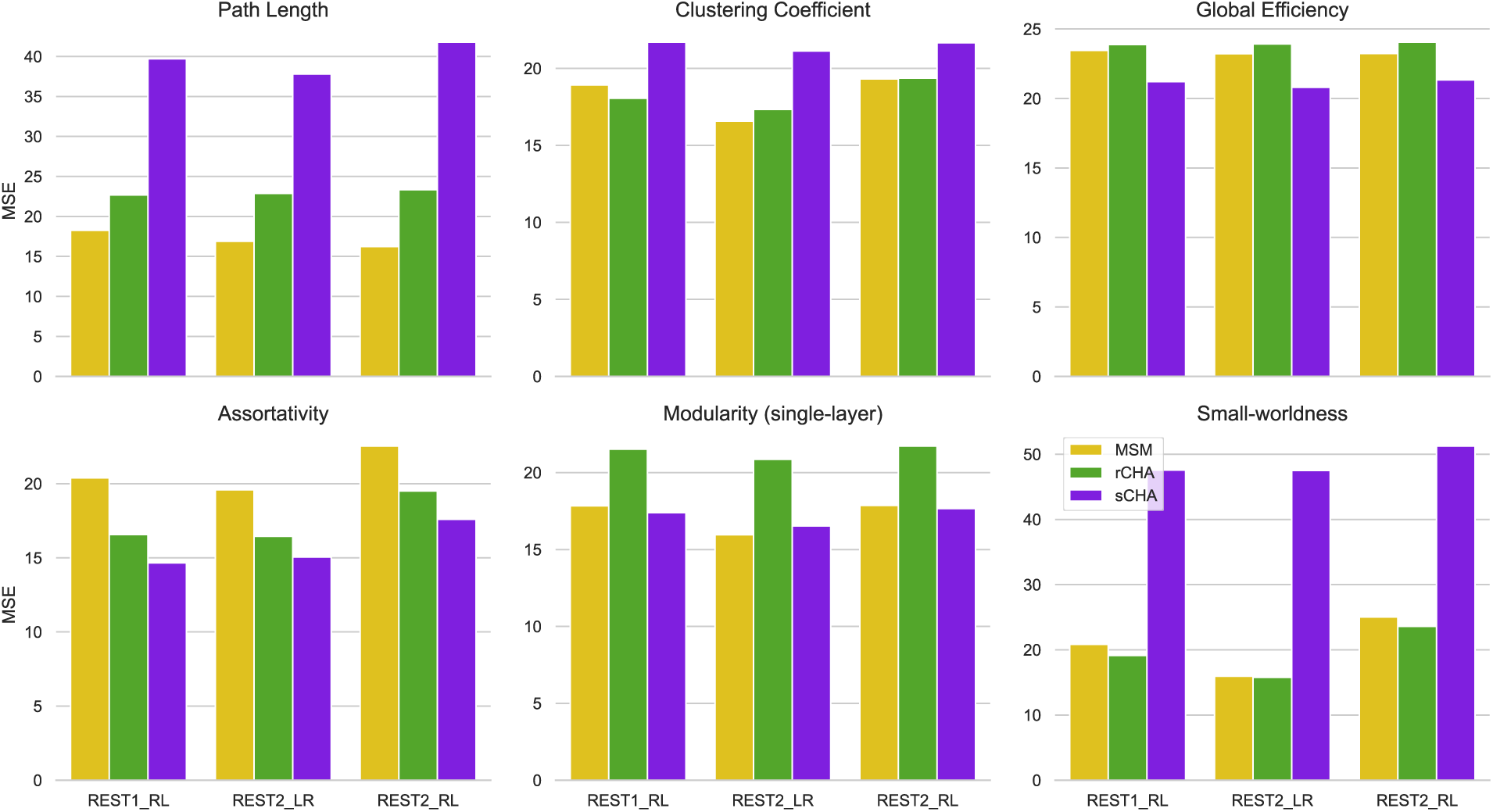
MSE evaluated before and after CHA for global measures in the three test datasets.

**Figure 6B** and **6C** visualize the MSE difference for the prediction of fluid intelligence between MSM and CHA aligned data (rCHA and sCHA, respectively) on the Glasser cortical map (lateral and medial views). This analysis incorporates measurements of eigenvalue centrality, closeness centrality, and clustering coefficient, which cover all three test sets. To facilitate comparison, we computed Δ/*max*(*absolute*(Δ)), where Δ is equal to *MSE_MSM_* – *MSE_CHA_*. This calculation yields a normalized value ranging from −1 and 1. Values closer to 1 and −1 indicate the superiority of CHA (depicted in a more bluish color) and MSM (depicted in a more reddish color) in predicting fluid intelligence, respectively. It is evident that most regions show positive values indicating lower MSE (i.e., improved prediction) after CHA, especially when considering closeness centrality and clustering coefficient. Regions that show lower MSE prior to hyperalignment correspond roughly to sensory regions of the brain, indicating that vertices vary less across individuals in these regions.

Additionally, we examined the average MSE across brain regions group by the 12 Cole/Anticivic networks, using both MSM and CHA data (refer to **Figure 6D**). This analysis allowed us to identify networks that demonstrated improved predictive capability for fluid intelligence following connectivity hyperalignment. Improvements were seen in all networks, except for the primary visual network when using sCHA data.

**Figure 8** shows the ridge regression results using global graph-theoretic patterns as features and compares the value of MSE in predicting fluid intelligence using various metrics before and after CHA, across all test sets. Notably, global efficiency and assortativity exhibit a decrease in MSE after sCHA across all three test datasets. This decrease in MSE implies that using a dataset highlighting an enhanced capacity for efficient information exchange and a more balanced arrangement of connections contributed to improved predictive performance. However, despite changes in observed network organization, no significant enhancement in prediction performance were found for path length, clustering coefficient, and small-worldness after implementing CHA. The results for single-layer modularity show inconsistent and nonsignificant changes across the three test data sets.

**Figure 8.**
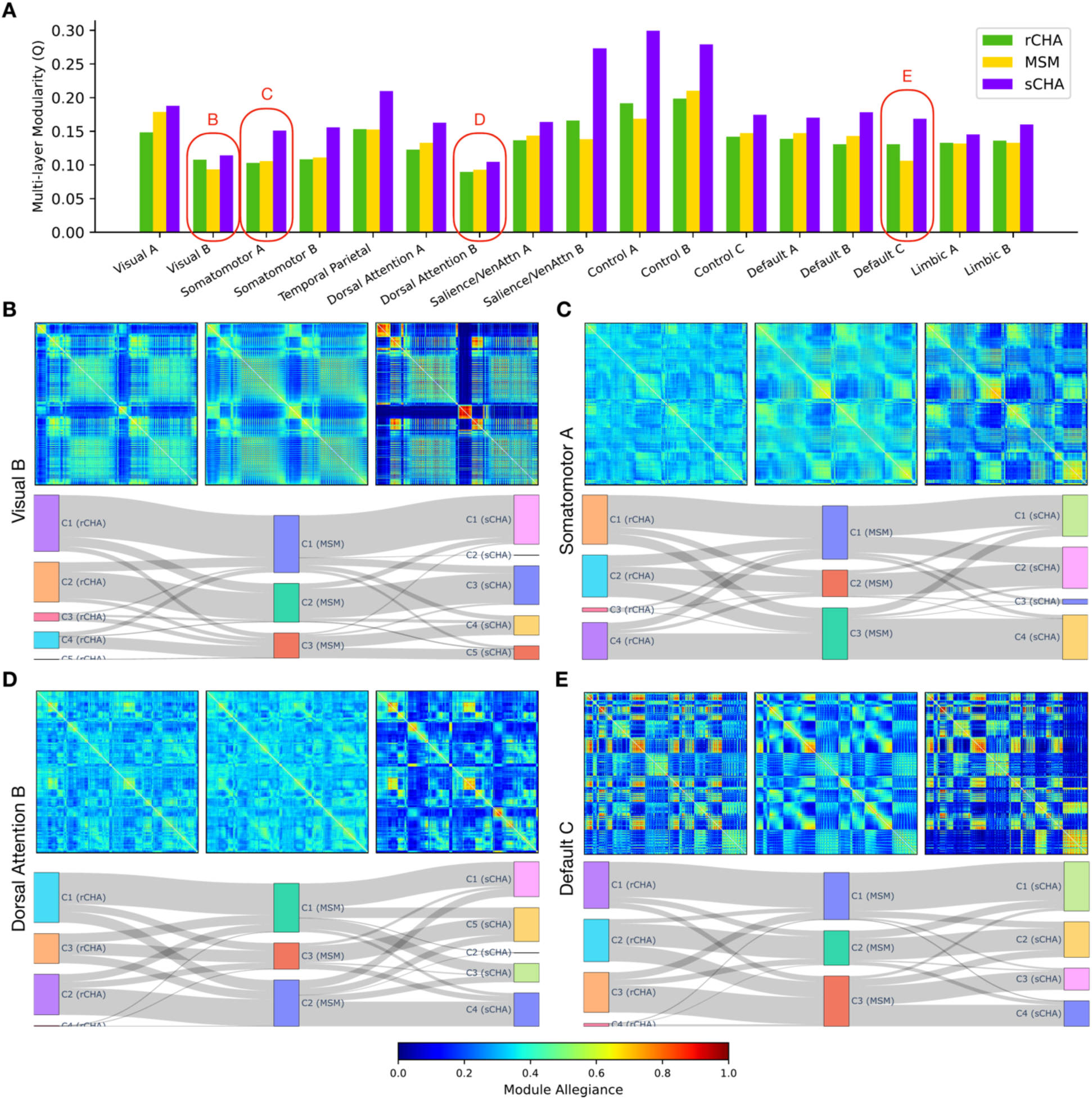
Multi-layer modularity properties before (MSM) and after connectivity hyperalignment (rCHA and sCHA). (A) Multi-layer modularity across 17 Yeo networks. (B-E, top) Module allegiance illustrating network engagement for the visual B, somatomotor A, dorsal attention B, and default C networks before (middle) and after rCHA and sCHA (left and right, respectively). (B-E, bottom) Detected communities for the aforementioned networks before and after connectivity hyperalignment and how they are intertwined (the middle panel is associated with the MSM, while the left and right panels corresponds to rCHA and sCHA, respectively).

### 3.4. Searchlight CHA improves module identification at the meso-scale

To probe the community structure of the fine-scale hyperaligned and non-hyperaligned data within each large-scale brain network, we employed a multi-layer (multi-subject) modularity framework (Mucha et al., 2010). Although early multi-layer studies began with uncovering patterns in temporal networks, we extended the notion to include the analysis of communities across individuals rather than time (Betzel et al., 2019; Zamani Esfahlani et al., 2021). As a result, each layer represents a subject’s connectivity matrix associated with other layers/subjects via an interlayer parameter to maintain consistent community labels across layers. The main outputs of the modularity maximization function include the modularity index, the number of communities/modules, and the community labels, which are discussed separately below. We performed the multi-layer community detection algorithm separately on the fine-scale connectivity matrices corresponding to each of the 17 Yeo networks.

**Figures 8A** shows the changes in modularity index *Q* before and after CHA for each of the 17 Yeo networks. Using a heuristic approach, we chose *γ* = 1.0 and *ω* = 0.1 for the structural resolution and inter-subject coupling parameters, respectively. **Figure 8A** illustrates that in all networks, the multi-layer modularity index, Q, increased after applying sCHA but not rCHA. These findings highlight an enhancement of meso-scale brain network community structure after sCHA. The higher modularity index indicates clearer functional groupings, likely improved functional specificity. The increased community count in hyperaligned data suggests finer functional subdivisions, implying a more detailed modular architecture. This collectively underscores how connectivity hyperalignment can uncover distinct functional units that hint at a stronger neural processing efficiency than previously thought.

The multi-layer modularity maximization algorithm produces community label (*C_ij_*) that assign modules to nodes in each layer (Bassett et al., 2015). These labels are utilized to construct allegiance matrices, as demonstrated in **Figures 8B**, **8C**, **8D**, and **8E** (top), which depict the allegiance patterns of four networks (visual B, somatomotor A, dorsal attention B, and default C networks) derived from the REST1_RL test set. For each panel/network, the top middle plot is associated with the MSM allegiance matrix, the top left is associated with rCHA, and the top right is associated with sCHA. These networks consist of varying numbers of vertices (3352, 5980, 4871, and 1489) distributed across different regions (35, 43, 83, and 39 regions) of the Glasser atlas. The module allegiance matrices provide insights into the engagement of brain vertices and their corresponding Glasser regions across subjects (Mattar et al., 2015). Refer to the Methods section for details on metric creation. As shown in **8B-E** (top), the reduction of inter-subject variations following sCHA resulted in more cohesive blocks, thus enhancing the identification of specialized modules within each network at a fine-scale. This improved ability to discern specialized communities following CHA likely contributes to the increased predictive power of fine-grained hyperaligned data, as demonstrated in the preceding section. **Figures 8B-E** (bottom) show the detected communities before and after hyperalignment. The middle panel is associated with MSM, while the left and right panels correspond to rCHA and sCHA, respectively, showcasing their relationships. These plots indicate a greater number of detected communities in hyperaligned data compared to non-hyperaligned data. Specifically, there are three detected communities for the MSM data and between four and five detected communities for both CHA data across these networks, with a more pronounced significance in sCHA.

Other informative properties, such as recruitment and integration, can be extracted from allegiance matrices to compare the modular structure of different populations (Bassett et al., 2015). For each network, these coefficients characterize the functional interplay between cortical vertices and predefined/static Glasser regions within that particular network. Recruitment evaluates how well a vertex is recruited to its predefined region, whereas integration evaluates the extent to which a vertex is integrated/connected with vertices in other predefined regions. For example, within the allegiance matrix of the default C network, each row (or column) represents a vertex whose recruitment and integration coefficients are determined by the average values inside and outside its static Glasser region, respectively. **Figures 8B-E** (top) show that cortical vertices are more likely to be recruited to their designated Glasser regions across subjects in sCHA than MSM and rCHA data, as demonstrated by the warm block-like patterns along the diagonal. In addition, the figure displays how, after sCHA, the integration of vertices between different regions has decreased in most parts of the network (cold blocks along the off-diagonal), while this coefficient has amplified between some regions (warm blocks along the off-diagonal), implying a possible significant and selective connectivity between them. In general, following sCHA, changes in recruitment and integration coefficients have resulted in a better identification of communities and their relationships within the network.

We extended the recruitment and integration analysis to other Yeo networks (17 in total), and the results are shown in **Figure 9**. **Figures 9A** and **9B** display the normalized recruitment/integration coefficients overlaid on brain cortical maps for rCHA and sCHA data, respectively, in comparison to MSM. They are computed as Δ/max(absolute(Δ)), where Δ represents the difference between Recruitment/Integration_MSM_ and Recruitment/Integration_CHA_. The distribution of recruitment and integration coefficients across cortical vertices are illustrated in **Figure 9C** and **9D**, revealing a substantial reduction in integration coefficients following sCHA. **Figure 9E** and **9F** show the mean recruitment/integration across each network’s vertices before and after CHA. Finally, **Figure 9G** and **9H** compare the integration coefficients for each network between the non-hyperaligned and hyperaligned data (rCHA and sCHA, respectively) as 17 different scatterplots. Accordingly, we found that vertices in MSM were more integrated with vertices of other regions than in CHA techniques.

**Figure 9.**
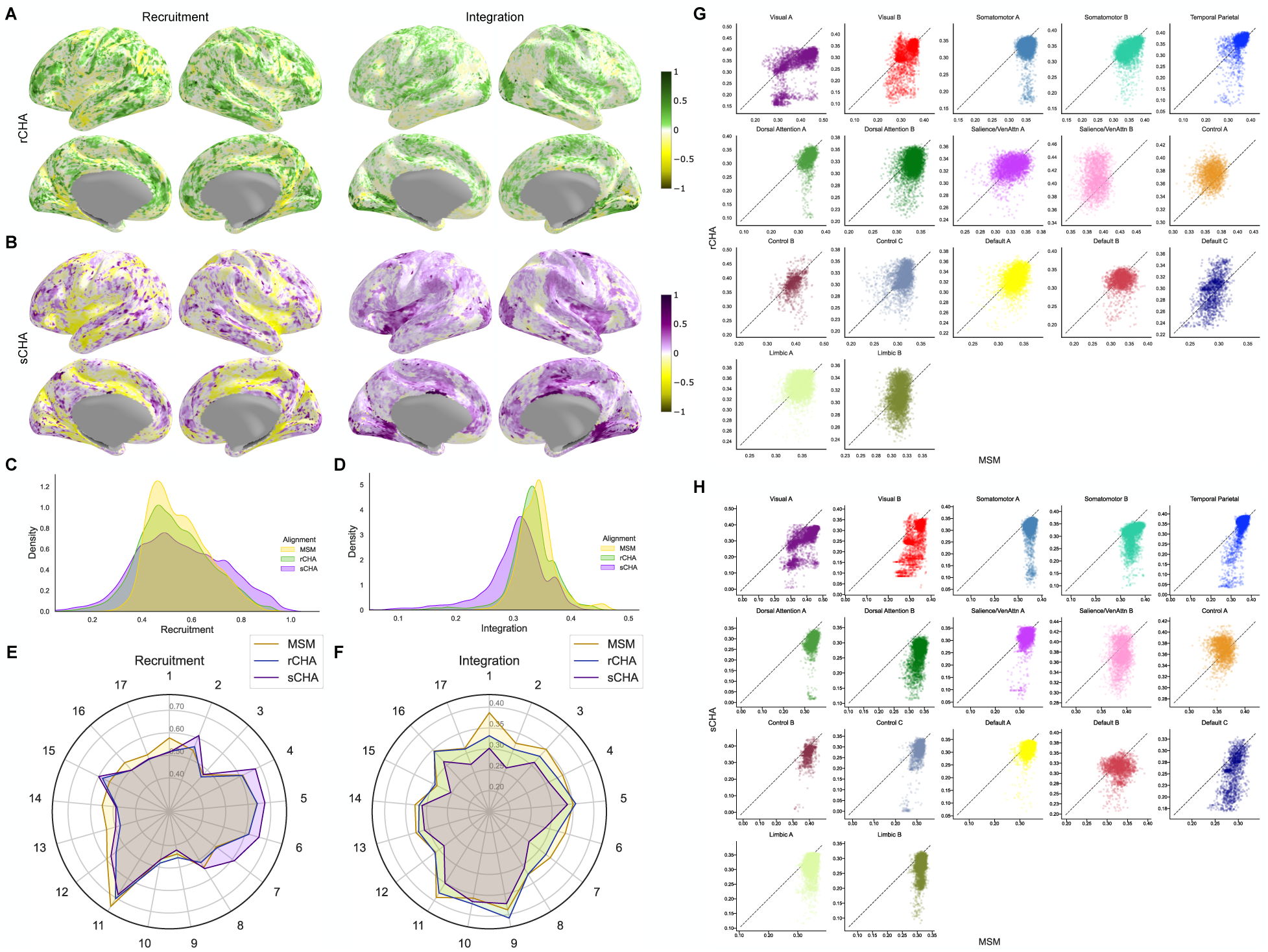
Recruitment and integration coefficients before and after CHA. Normalized coefficients between (A) MSM and rCHA, and between (B) MSM and sCHA. (C, D) Distribution across cortical vertices. (E, F) Average across network vertices for recruitment and integration coefficients pre- and post-CHA. (G, H) Integration coefficient comparison for 17 Yeo networks.

## 4. Discussion

In this paper, we demonstrate how CHA (including both rCHA and sCHA) improves individual-specific information and enhances the performance of human connectome graph analysis at both fine and coarse spatial scales. Fine-scale structure refers to spatial granularity at the voxel or vertex level, in contrast to the coarser structure defined by brain parcels or regions (i.e., sets of voxels). In CHA, fine-scale local variations in connectivity profile are captured using a set of shared basis functions across subjects within a high-dimensional information space. The introduction and advancement of CHA has opened up an untapped territory in connectome analysis, enabling in-depth exploration of fine-scale brain structure (Guntupalli et al., 2018). For instance, recent work by Feilong et al. (2021) using rCHA showcased superior predictive power of fine-scale connectivity patterns over coarser-grained counterparts for general intelligence prediction. Our work brings hyperalignment for the first time into the realm of network neuroscience, enabling researchers to scrutinize brain network topology with higher spatial resolution – down to individual voxels or vertices – a level of detail unattainable with traditional coarse or smoothed data approaches. This study has two primary objectives: investigating CHA’s impact on the observed brain topological properties across both coarse and fine scales, and exploring whether this functional alignment leads to enhanced prediction of fluid intelligence using fine-scale graph theory patterns.

The coarse-scale analysis of graph metrics following CHA compared to MSM alignment yields insightful observations about the network properties and their potential implications. Notably, the shift towards decreased global efficiency coupled with increased clustering coefficient, assortativity, single-layer modularity, and small-worldness metrics post-CHA implies that brain organization may be more localized and interconnected than previously thought. This paradoxical decrease in global efficiency might signify a trade-off between network-wide information transfer and the establishment of specialized communication hubs, aligning with the brain’s efficient processing through specialized networks. The rise in network attributes supports the notion of a “small-world” architecture, where interconnected modules optimize the balance between segregation and integration.

In exploring local properties at coarse-scale, CHA uncovers shifts in centrality metrics and nodal clustering coefficients within distinct brain networks, redefining hubness and local connectivity. Enhanced centrality and increased nodal clustering in networks like secondary visual, somatomotor, cingulo-opercular, dorsal attention, and posterior multimodal suggest improved local integration and information flow efficiency. However, when applying these topological patterns to predict fluid intelligence, sCHA doesn’t show increased accuracy. This suggests that while CHA alters network connectivity measures, it may not improve the ability to capture inter-individual differences or external validity at the course scale. In essence, we hypothesize that smoothing hyperaligned fMRI data across coarse-grained parcellations could lead to the removal of any remaining idiosyncrasies.

An important question arises as to why, even with the coarse-scale brain parcellation (i.e., averaging fine-grained data across regions), significant topological differences persist within the brain network after the implementation of sCHA. Does this regional averaging not counteract the impact of alterations in cortical topography of the information that occur at local fields (i.e., searchlights)? The response lies in the characteristic of searchlight HA (Guntupalli et al., 2016), as distinguished from ROI HA (Haxby et al., 2011), where information can flow both into and out of a brain region, particularly near boundaries. This design feature ensures that the same function does not consistently reside in a certain anatomical region across subjects (Eickhoff et al., 2018; Feilong et al., 2021). This dynamic flow of information, encompassing noise as well, potentially constitutes the neural basis for the observed topological changes resulting from CHA at the coarse scale.

In contrast to the coarse-grained counterpart, our fine-grained results showed that local features extracted from CHA-aligned data exhibited superior performance in predicting intelligence compared to MSM-aligned data, indicating an improvement in discerning topographic idiosyncrasies at finer granularity. Local features primarily convey information about the centrality of the nodes or the interconnections among the neighbors of the nodes, which are sometimes correlated with each other. The fine-grained results additionally underscored that the most effective predictors of intelligence reside in the orbito-affective, auditory, cingulo-opercular, ventral multimodal, somatomotor, and default mode networks, respectively. Collectively, our findings suggest that the application of CHA in fine-grained analysis represents an important new processing step in brain network research. This approach enables the capture of meaningful performance-related activity at a heightened spatial granularity, bypassing the need for smoothing. Signals at this scene have previously been overlooked due to factors such as averaging and smoothing procedures designed for enhancing consistency in group analyses of fundamental task-related activity, assumptions that such activity constitutes noise, signal dropout, and computational cost.

We expanded our investigation by performing a multi-layer (multi-subject) modularity analysis on each brain network separately using fine-grained data, examining the changes in network organization before and after CHA. This mesoscale analysis enables the exploration of how different regions/vertices are interconnected, providing a direct measure of network organization using module allegiance matrix, recruitment, and integration coefficients. Examination of allegiance matrices derived from non-hyperaligned data shows that numerous connections between brain regions and vertices in each network are erroneously estimated due to misalignment. Put more simply, without CHA, discrete modules are blurred together. These erroneous connections were vastly reduced after CHA, resulting in better module identification. This modular refinement aligns with our fine-grained results, wherein local features extracted from CHA-aligned data demonstrated superior predictive performance in fluid intelligence compared to MSM-aligned data. This signifies an elevated capacity to discern topographic idiosyncrasies at a more refined scale.

Our meso-scale analysis also revealed a consistent decrease in the integration coefficient—a metric that assesses the degree of vertex integration with other regions across individuals/layers—across all brain networks after applying CHA, indicating a more structured network generated by hyperaligned data. This underscores CHA’s impact on network organization. The reduction of erroneous connections, heightened module identification, and emergence of sub-modules collectively highlight CHA’s efficacy in enhancing inter-region connectivity estimation precision. This modular refinement confirms CHA’s ability to form a coherent brain connectivity representation, supporting improved predictive performance of fluid intelligence through finer-scale local features.

To conclude, this study not only contributes valuable insights into the impact of CHA on brain network properties at varying spatial scales but also underscores the potential for fine-grained analysis using this approach to reveal previously unexplored facets of brain structure and function. The observed effects on global, local, and mesoscale network properties provide a nuanced understanding of how CHA influences observed brain connectivity and organization. These findings hold promise for enhancing our comprehension of individual differences, cognitive processes, and brain disorders, with implications spanning from basic neuroscience to clinical applications.

### Future directions and limitations

Certain limitations associated with this study should be considered in future research. First, the inter-regional connections of the cortical vertices are not considered during fine-grained network investigation. Here we built a network of nodal interactions within each Glasser region and examined them separately, which does not cover all functional connectivity within the whole cortex. Considering whole-brain connections at fine spatial scales (i.e., both intra-regional and inter-regional links) is computationally expensive and beyond the scope of the current work. One direction for future study could be how to use maximum brain information while tackling computational complexity.

Another issue concerns the searchlight implementation. Since the location of a functional area and its boundaries is not the same across individuals, the searchlight approach is generally preferred to the ROI-based approach in HA studies (Feilong et al., 2021). However, the main weakness of this method is increased difficulty in interpreting the results. For example, an improvement in prediction performance after searchlight hyperalignment can be caused by better alignment of functional representations within a region. Still, it can also be caused by the inclusion of additional information from neighboring functional regions, as the searchlight may cross area boundaries.

Finally, another possible extension of this work includes studying the HCP task fMRI dataset to derive a common hyperalignment model (i.e., functional space) and transformations, then analyzing a broader set of task fMRI data by mapping it into this space. It is possible to train hyperalignment using one dataset and apply it to another, as long as data are separated into independent training (HA mapping) and test data sets. Thus, a shared, large-sample common space could facilitate future predictive models across many datasets.

## Data and code availability statement

Data were provided by the Human Connectome Project, WU-Minn Consortium (Principal Investigators: David Van Essen and Kamil Ugurbil; 1U54MH091657) funded by the 16 NIH Institutes and Centers that support the NIH Blueprint for Neuroscience Research, and by the McDonnell Center for Systems Neuroscience at Washington University. Python code for implementing our analyses and visualizations can be found via the following link: https://github.com/fvfarahani/hyperaligned-brain-network in which we used different packages such as NumPy, Pandas, SciPy, hcp-utils, Nilearn, Teneto, Matplotlib, and Seaborn. The code for performing searchlight hyperalignment was adapted from PyMVPA (Hanke et al., 2009; http://www.pymvpa.org/). Graph theory measures were calculated using the Brain Connectivity Toolbox (https://sites.google.com/site/bctnet/). The GenLouvain MATLAB package (Jutla et al., 2011) used for multi-layer community detection is freely accessible online (https://github.com/GenLouvain/GenLouvain).

## CRediT authorship contribution statement

**Farzad V. Farahani:** Methodology, Formal analysis, Writing - Original Draft, Visualization, Validation; **Mary Beth Nebel:** Writing - Review & Editing; **Tor D. Wager:** Writing - Review & Editing; **Martin A. Lindquist:** Methodology, Funding acquisition, Writing - Review & Editing, Supervision.

## Footnotes

1 The modularity matrix, *Bij*, is a full matrix whose entries represent the difference between observed (*Aij*) and expected (*Vij*) connection weights (B = A - V).

